# GreenMaps: a Tool for Addressing the Wallacean Shortfall in the Global Distribution of Plants

**DOI:** 10.1101/2020.02.21.960161

**Authors:** Barnabas H. Daru

## Abstract

The exponential growth of species occurrence data can facilitate dynamic biodiversity analyses. However, raw biodiversity data alone should not be used indiscriminately due to inherent sampling biases, impediments that contribute to Wallacean shortfall (i.e. the paucity of species’ geographic information). It has been suggested that Wallacean shortfall is a common phenomenon across taxa, however, there is no global assessment geared toward overcoming this impediment for plants, despite the fundamental role of plants in ecosystem stability, food security and biodiversity conservation. Here, I present GreenMaps, a new tool that will permit a rapid initial assessment of the Wallacean shortfall for plants by building base maps of species’ predicted distributions upon which citizen science participation could contribute to spatial
validation of the actual range occupied by species. The initial stages of GreenMaps have now been accomplished, providing a massive dataset of modeled range maps for over 194,000 vascular plant species. This will make it the largest and only global assessment of geographic distributions for plant species at scales relevant to research and conservation. Ultimately, GreenMaps will interface with a mobile application to enable volunteers from any region of the world to validate predicted species distributions to be used for the generation of new and improved global map of plant distributions.

## 1.0 Shortfalls in Knowledge of the Geography of Plants

Understanding patterns, underlying processes and principles governing biodiversity at large scales requires an accurate knowledge of species geographic distributions. The increasing availability of biodiversity data from multiple heterogeneous sources, including museum collections, observation records, biological literature and local databases, has led to their compilation into major data hubs such as the Global Biodiversity Information Facility (GBIF; Edwards et al. 2000), iNaturalist (iNaturalist 2019), and iDigBio (www.idigbio.org), that can facilitate macroecological analyses (Díaz et al. 2016; Pollock et al. 2017; Zanne 2018; Daru et al. 2019). These datasets are often point occurrences of where a species has been documented as present based on a voucher specimen in a museum/herbarium or reliable sighting in the field. However, the raw occurrence data alone should not be used indiscriminately due to prevailing sampling biases, gaps and uncertainties inherent in this type of data (Lira-Noriega et al. 2007; Meyer et al. 2016; Stropp et al. 2016; Daru et al. 2018). Another disadvantage of raw occurrence data is that they do not extrapolate beyond relatively few known locations to make inferences about the potential geographic range occupied by a species. Such impediment is often called Wallacean shortfall, the paucity of information on species distributions at a global scale (Hortal et al. 2015), is more acute for less-charismatic groups such as invertebrates and plants, and can induce strong limitations in conservation and spatial planning (Hortal et al. 2008; Boakes et al. 2010). For example, existing plant occurrence information is heavily skewed towards better sampled or more accessible areas and regions where digitisation is more advanced (e.g. North America, Western Europe), whereas some of the world’s most biodiverse regions, for example, Central Asia, Africa or Amazonia, remain heavily under-represented in botanical sampling (Bush and Lovejoy 2007; Meyer et al. 2016). Progress in addressing some of the key questions in biodiversity science, such as how diversity is distributed in space, how it changes across time, and how it contributes to the resilience of ecosystems to global change, depends heavily on our understanding of accurate species distributions. It is therefore critical to address the Wallacean shortfall in species distributions by improving the global coverage and representativeness of biodiversity data.

In order to reduce the Wallacean shortfall in species distributions, there have been recent efforts incorporating niche-based models for a variety of taxonomic groups. Examples include efforts from Aquamaps (Kaschner et al. 2013) for marine species, and Lifemapper (Beach et al. 2015) and Map of Life (MoL, http://www.mol.org; Jetz et al. 2012) for terrestrial vertebrates. Because these initiatives complemented ecological data (e.g. climate and functional traits) with fine-scale aggregated data (e.g. specimen records, surveys, and experiments) to model species distributions, they create strong synergies that bridge the gap between fine-scale precision and global representativeness (König et al. 2019). So far, however, there is no solution that incorporates modeled distributions and online mobilisation for plants at a global scale in a generic way. Plants are foundation species, sustaining food chains and driving terrestrial ecosystem productivity (Loreau et al. 2001; Zak et al. 2003). Consequently, plant distribution often underlies the biogeographic histories of other organisms, and thus a global distribution map for plants is critical for predicting and conserving diversity at other trophic levels.

Vascular plants form a very diverse taxonomic group comprising about 300,750 species worldwide (Christenhusz and Chase 2014; APG IV 2016; Christenhusz et al. 2017). Plants occur across all types of biomes, from rainforests to savannas. Some are cosmopolitan, spanning borders e.g. the Cogon grass (*Imperata cylindrica*), whereas some species occur in very restricted distributions of a few hectares such as the Olympic violet (*Viola flettii*) endemic to the Olympic Mountains of Washington in United States. The overall ecosystem services provided by plants are estimated to be worth at least US$16–54 trillion per year worldwide (Costanza et al. 2007). These services include provisioning (e.g. food and medicines), regulation of ecosystem processes (e.g. trophic regulation, water purification), cultural (firewood, ornamental) and supporting services (e.g. primary productivity), but yet estimates of plant geographic distributions at a global scale rest largely on extrapolations. Species distribution models (SDMs) provide an unbiased and easily interpretable estimate of improving representativeness and coverage of species distributions (Peterson et al. 2011).

To date, studies utilising SDMs to improve coverage of plant occurrences have focused mostly at the regional or country scale despite species distributions spanning political and regional borders, calling for a truly global assessment of species distributions (Jetz et al. 2019). To accurately depict the geographic range for every vascular plant species represents great challenges in terms of computation and global coverage at the species level. For instance, an examination of GBIF reveals that available point occurrences covered most plant species at only seven or fewer unique locations (Meyer et al. 2016), too few for estimating robust species distribution models (Guisan et al. 2007; Feeley and Silman 2011) or generating geographic range maps (Gaston and Fuller 2009; Rivers et al. 2011), and thus might require additional filtering, human labor and continuous update. I proposed to use climate as reliable predictor of plant distributions at broad scales, consistent with Hutchinson’s multidimensional niche concept that every species is limited by a number of environmental factors (Hutchinson 1957), coupled with well-established relationships between plant species richness and climate (Currie 1991; Olson et al. 2001; Wiens and Donoghue 2004; Fine 2015). Thus, I define the probability of occurrence of a species as the set of favourable environmental conditions that allow a species to survive and reproduce, and populations to maintain their numbers (**Fig. 1**). A global analysis of this nature should identify common links between species’ ecological preferences and potential biomes that are most susceptible to changes in community composition under global change (Hooper et al. 2005; Cadotte et al. 2009; Barnosky et al. 2012; Lewis and Maslin 2015).

**Fig. 1.**
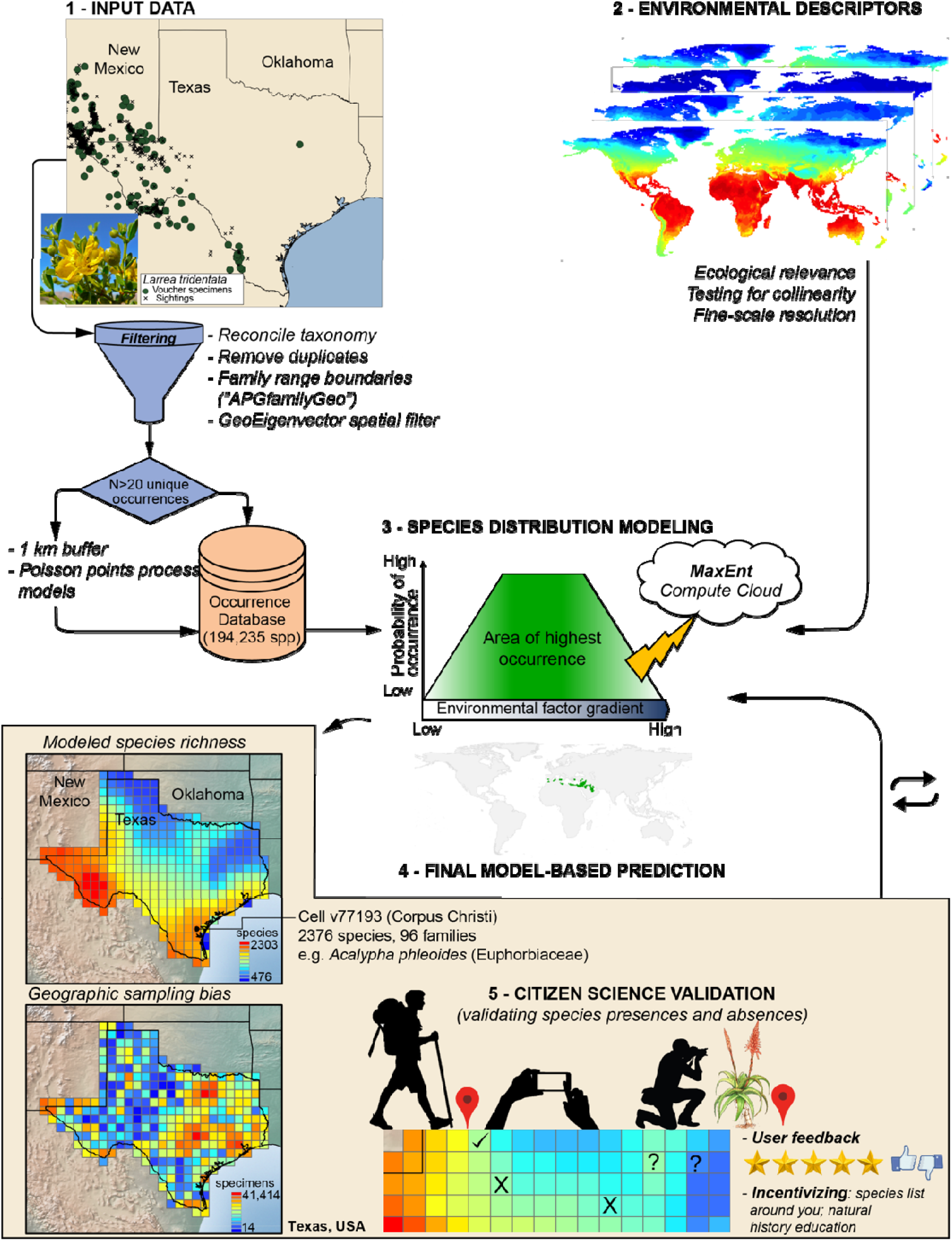
Overview of the workflow of GreenMaps showing archive data and citizen-defined maps. Products from the first phase of GreenMaps (1 to 4) have been accomplished, providing model-based predictions for 194,235 species. Phase 2 will involve citizen science validation of the species predictions.

Here, I develop GreenMaps, a novel cyberinfrastructure that permits a rapid initial assessment of the Wallacean shortfall for plants by building base maps of species’ predicted distributions based on SDMs upon which citizen science participation could contribute to spatial validation of the actual range occupied by species. Ultimately, GreenMaps will interface with a mobile application to enable volunteers from any region of the world to validate predicted species distributions to be used for the generation of new and improved global map of plant distributions at scales relevant to research and decision-making. Specifically, I address three objectives. 1) Design and development of GreenMaps, toward addressing the Wallacean shortfall in plant distribution. These requires high performance computing strategies for species distribution modeling spanning 300,000 species of vascular plants at a global scale. 2) Design and development of easy to use mobile application for GreenMaps to be used by biodiversity scientists and citizen scientists alike for spatial validation of the actual range occupied by species. 3) Assess how plant diversity is distributed in space, and explore the tempo and mode of plant diversification on mountains.

## 2.0 Design and Development of GreenMaps: How it Works

GreenMaps consists of two primary types of data - GreenMaps archive data and citizen-defined maps (**Fig. 1**). GreenMaps archive data consists of global model-based species distributions along with occurrence records and carefully vetted taxonomic names (species, genus, family, order and clade) for all vascular plants at a global scale. Citizen-defined maps consist of an interface in which experts and citizen scientists can review, edit and approve these predicted range maps. The main features of the tool are described below.

In the first stage of GreenMaps, I generated the base maps of predicted species’ distributions for citizen science validation. To this end, I obtained occurrence records from specimen records, online repositories (e.g. iNaturalist, GBIF, iDigBio, records from the primary literature, and personal observations). These records were thoroughly cleaned to reconcile old names/synonyms to current taxon concepts e.g. The Plant List (www.theplantlist.org), and/or to remove duplicates and records with doubtful or imprecise localities (e.g. points in the sea and records with inverted coordinates). Applying such cleaning filters wiped out most available occurrence records, resulting in too few records for the vast majority of plant species. To improve the utility of species with too few records for SDMs, I created buffers of 1-2 km radius as spatial offsets in Poisson point process model around each point (see Merow et al. 2016) and randomly (Poisson) distributed points (adding up to 20) within the boundaries of the buffers (a good SDM requires 10-200 unique records) (Kadmon et al. 2003; Rivers et al. 2011). These points were used in addition to the cleaned dataset as inputs for the species distribution modelling (**Fig. 1**).

In a second step, I imposed two stringent spatial filters to restrict the area of species to their known native ranges and to prevent erroneous records and predictions in areas that might contain suitable habitat but are not occupied by a species. *1) APGfamilyGeo:* I bounded every species distributions based on expert drawn occurrence polygons (“expert maps”) of global plant family geographic distributions (Heywood 1993; APG IV 2016). Although expert maps have limitations such as encompassing areas of unsuitable habitat within the range boundaries (Merow et al. 2017), they ultimately represent excellent resources for delimiting the boundaries beyond which a species is not expected to occur. This spatial filter was applied to constrain species predictions during the modeling process. *2) GeoEigenvector:* These are orthogonal variables representing spatial relationships among cells in a grid, encompassing the geometry of the study area at various scales (Diniz-Filho et al. 2005). The software SAM (Spatial Analysis in Macroecology; Rangel et al. 2010) was used to generate a pairwise geographical connectivity matrix among grid cells as input to establish a truncation distance for the eigenvector-based spatial filtering, resulting in a total of 150 spatial filters. These filters were then resampled to the same resolution as the environmental variables and were included with the bioclimatic variables for the SDM.

Third, I combined the occurrence records with 19 georeferenced bioclimatic variables at a resolution of 30 arc-second from WorldClim (Hijmans et al. 2005). These variables correlate with key physical and biological attributes of many plant species at a scale close to *in situ* records (Jetz et al. 2019) such as precipitation, temperature, primary productivity, plant functional traits, elevation (Hansen et al. 2013; Tuanmu and Jetz 2015; Jetz et al. 2016; Wilson and Jetz 2016), and sometimes the actual niche realised by species directly (Fretwell and Trathan 2009; He et al. 2015; Asner and Martin 2016). Environmental data can also be derived from MODIS, LandSat, NLCD, PRISM, and EarthEnv.

The algorithm MaxEnt (Phillips et al. 2006) was used as the basis for modeling the distributions of plants not only because it works with presence-only data but it is effective and sensitive to differences in modelling settings and can accommodate large datasets that span regional and global scales (Shcheglovitova and Anderson 2013; Syfert et al. 2013; Warren and Seifert 2011). MaxEnt estimates a species’ probability distribution that has maximum entropy, subject to a set of constraints based on knowledge of environmental conditions at the known occurrence locations (Phillips et al. 2006). I integrated cloud-based statistical workflows via the High Performance Computing Clusters of Texas A&M University-Corpus Christi (https://hpc.tamucc.edu/) implemented in R (R Core Team 2018) and modeled the potential distribution of 194,235 terrestrial vascular plant species using the presence only data and 10,000 background points. I used the equal training sensitivity and specificity threshold (Liu et al. 2005) to create presence-absence maps. Model performances were evaluated using the Area Under the Receiver Operator Curve (AUC; Bewick et al. 2004). The final outputs from MaxEnt consisted of a modeled range map (stored in raster format at grid cell resolution of 0.5 degree equivalent to 50 km) and the observed point occurrences (**Fig. 1**). A modeled range map resembles a species’ fundamental niche, its environmental suitability, its likelihood of being collected, and its native range area of occupancy as known from its family’s expert drawn geographic range (Heywood 1993; APG IV 2016).

A total of 194,235 modeled range maps, representing 194,235 species and 222 families have been generated and are available at a resolution of 0.5° × 0.5° (**Fig. 2**), making it the largest and only global assessment of geographic distributions for plants at the species-level. Based on these base maps alone, I found that the geographic distribution of plant species is very uneven, with higher diversity in the tropics peaking in Amazonia, Andes, southern Africa, Madagascar, Himalaya-Hengduan mountains, Southeast Asia, Southern Europe and Central America, mirroring global latitudinal diversity gradient (**Fig. 2**). My predicted distributions contrast sharply to observed patterns of sampling biases in plant occurrence records (Meyer et al. 2016), lending strong support and power to the modeled distributions. For example, my modeled maps highlight well-known plant diversity hotspots that have essentially no virtual collections available e.g. Amazonia and Himalaya-Hengduan Mountains. On the other hand, a point map (stored in CSV format), displays the locations of the georeferenced occurrence of a species following data cleaning. A third layer of GreenMaps archive data corresponds to map of geographic biases in botanical sampling density. Here, I used the observed occurrence points to map geographic biases in sampling density (**Fig. 1**), defined as grid cells of excessive (hotspots) or insufficient (coldspots) collection (Hijmans et al. 2000; Daru et al. 2018). The initial milestones of GreenMaps have now been implemented and include modeled range maps, point maps and bias maps for 194,235 species (222 families) and will be stored on a web portal as base maps for citizen science validation. Code for running the species distribution models, mapping hotspots and coldspots of sampling density, generating species richness, and endemism is publicly available in a new R package *bioregion* (Daru and Schliep 2019).

**Fig. 2.**
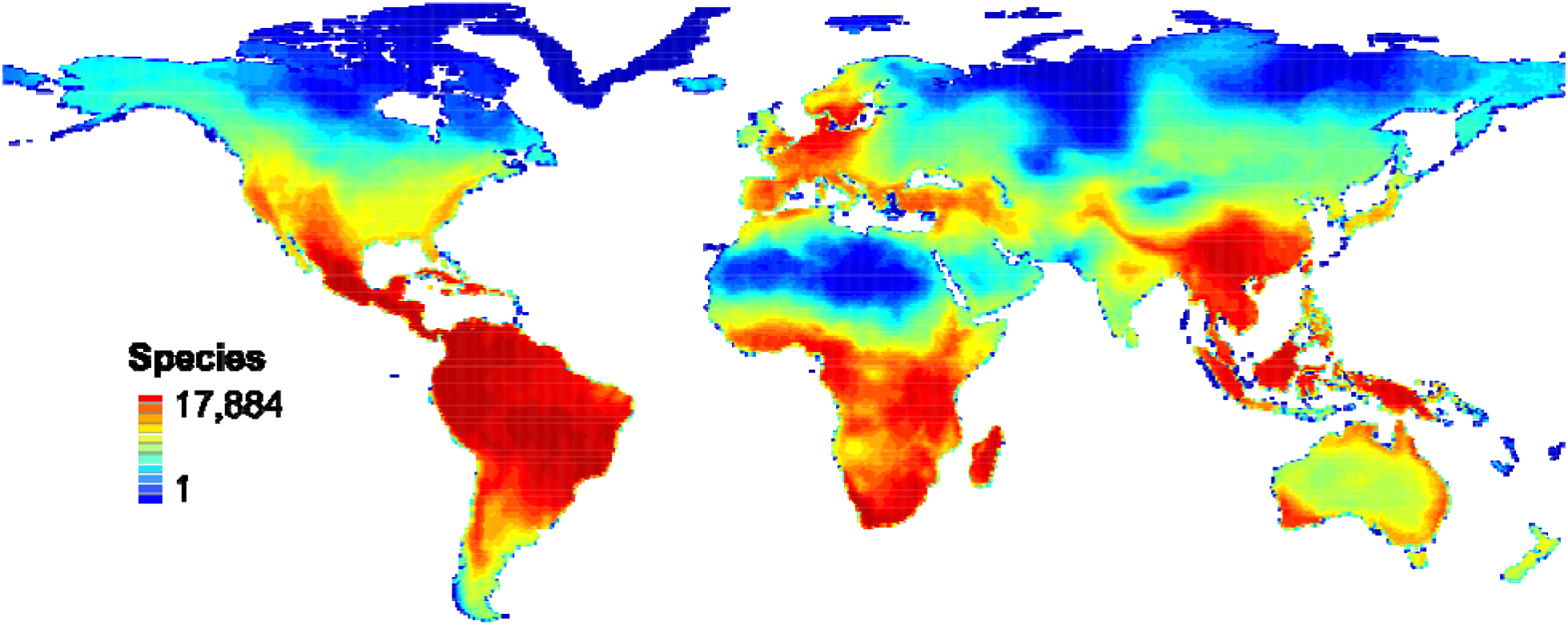
The geographic distributions of global patterns of vascular plant species richness. The map shows predicted distributions for 194,235 species (221 families) based on species distribution models within grid cells at a resolution of 0.5° × 0.5°, ranging from deep blue (minimum 1 species) to deep red (maximum 17,884 species).

### 2.1 Next Steps for GreenMaps

Two major steps are required for the full implementation of GreenMaps: 1) generating additional modeled base maps for the missing 106,515 species (out of the estimated 300,750 species; Christenhusz and Chase 2014; APG IV 2016; Christenhusz et al. 2017). I expect that the majority of the missing species will likely already be rare, since a local distribution is one of the major determinants in their escaping detection so far; and 2) the implementation of citizen-defined maps to ensure that the modeled range maps are technically valid and representative of the species native range area. Because the raw point occurrences for most plant species are too few for estimating robust species distribution models or generating geographic range maps (Guisan et al. 2007; Feeley and Silman 2011), citizen science participation is a relatively inexpensive form of contributing data to overcome the gaps in botanical sampling. Conceptually, the accuracy of a good SDM depends on the quality of the input occurrence records and the area restrictions imposed for a species (Phillips et al. 2006). Thus, GreenMaps provides volunteers access to an interactive mode that allows them to edit maps via a mobile app or web browser. In this mode, volunteers can exclusively choose species of interest based on specific traits and change the mapping parameters using pencil and eraser tools. For example, volunteers can explore which trees are common in an area, which species are succulent, herbs, or epiphytes; or by taxonomic group such as ferns, gymnosperms, monocots, orchids, grasses etc. Since almost any information can nowadays be geoTagged with appropriate geographic location (Goodchild 2007; Sui 2008), I implemented the option that allows volunteers to collect ancillary data such as pictures, recordings, and DNA sequences, upload them and metadata will be populated automatically.

Another promising feature of GreenMaps is that users can validate the occurrence of a species from any 0.5° × 0.5° pixel on the globe by voting whether or not a species is actually present or absent in the area as predicted by the modeled distributions. For instance, GreenMaps predicts the occurrence of 6118 plant species (representing 125 families, 39 orders) in the state of Texas, USA, with areas of high predicted plant species richness in the southwest running from Nuevo Laredo in Mexico to Big Bend National Park Texas (**Fig. 3a**). When I weighted each grid cell by the number of species actually observed in the State, I found a mismatch between areas of high predicted richness versus observed richness, suggesting that the sampling of plant species in Texas is non-random, but clustered around big cities such as Austin, San Antonio, Houston, Dallas and Corpus Christi (**Fig. 3b**). Indeed, only 25.5% (2023 out of 7924) of observed species distributions overlap the geographic ranges of species modeled by GreenMaps (**Fig. 3c**). This overlap increased to 67.7% when assessed at the family level, with 12-42% of families unique to either modeled distributions or observations (**Fig. 3d**). Thus, more observations are required in places that are under-sampled, but predicted to have more diversity. Using rarefaction, the observed occurrences did not fully predict plant species diversity in Texas (**Fig. 3e**), further highlighting inherent sampling biases in plant occurrence records (Meyer et al. 2016; Daru et al. 2018). I also found a tendency of the observed occurrences to concentrate in particular clades as opposed to more dispersed phylogenetic distributions for the modeled distributions (**Fig. 4**), lending support to the utility of species distribution models in overcoming impediments of sampling biases. Thus, GreenMaps will be designed by implementing these features (observed sampling density vs predicted distributions, phylogenetic representation) to realize a truly robust assessment of plant diversity at scales relevant to research.

**Fig. 3.**
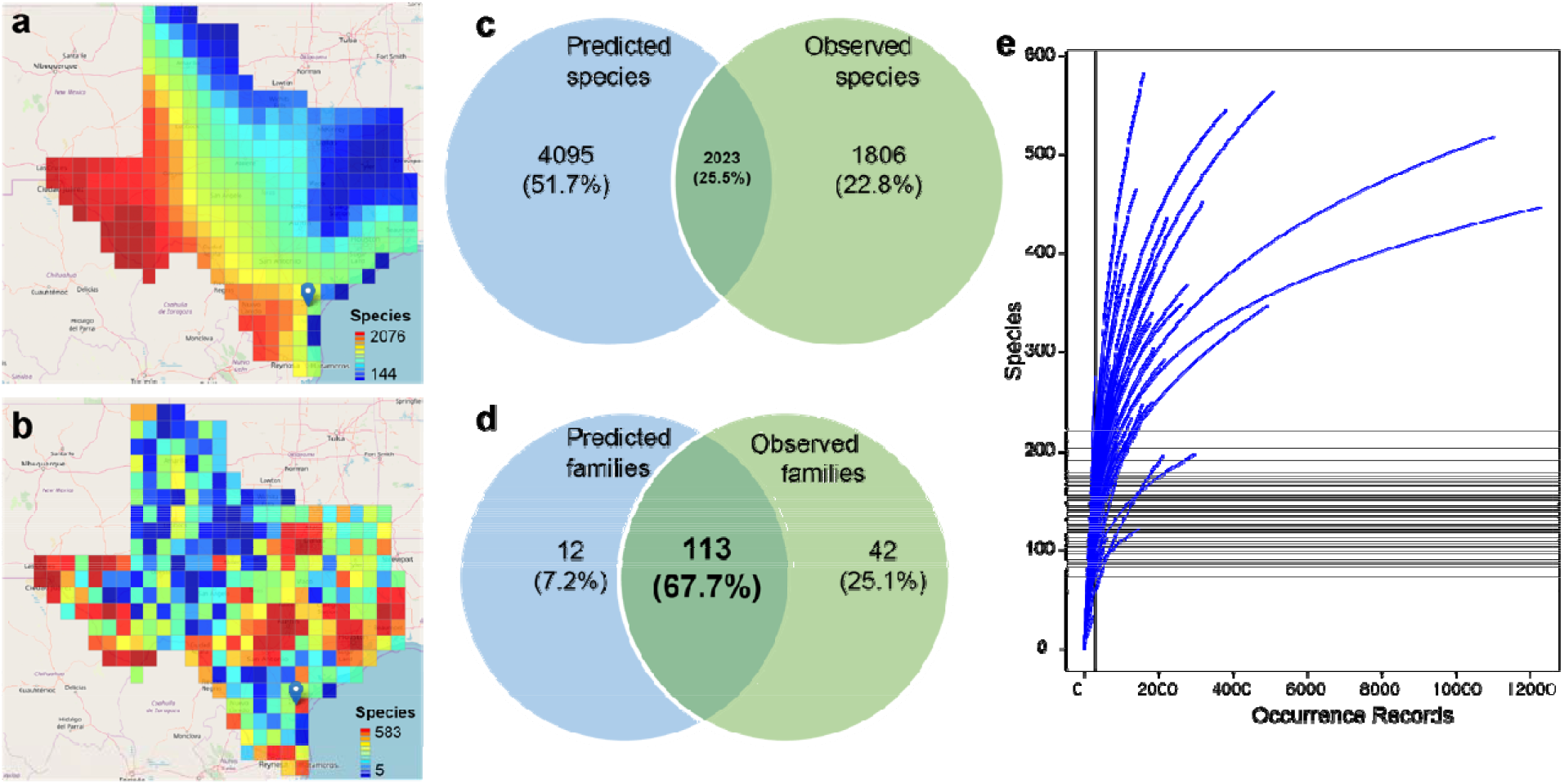
Overlap and mismatch between predicted and observed distributions of plants in Texas, USA. **a,** Predicted species richness (n = 6118 species). **b**, Observed species richness (n = 3829 species). **c**, Venn diagram of overlap and incongruence in the number of species predicted versus observed (predicted species ∪ observed species = 7924 species). **d**, Overlap and incongruence in the number of families predicted to occur versus families actually observed in Texas USA (predicted families ∪ observed families = 167 families). **e**, Observed occurrence-based rarefaction sampling curves (blue lines) for species richness for the vascular plants of Texas, USA. Each blue line represents the rarefaction curve for different locations within Texas.

**Fig. 4.**
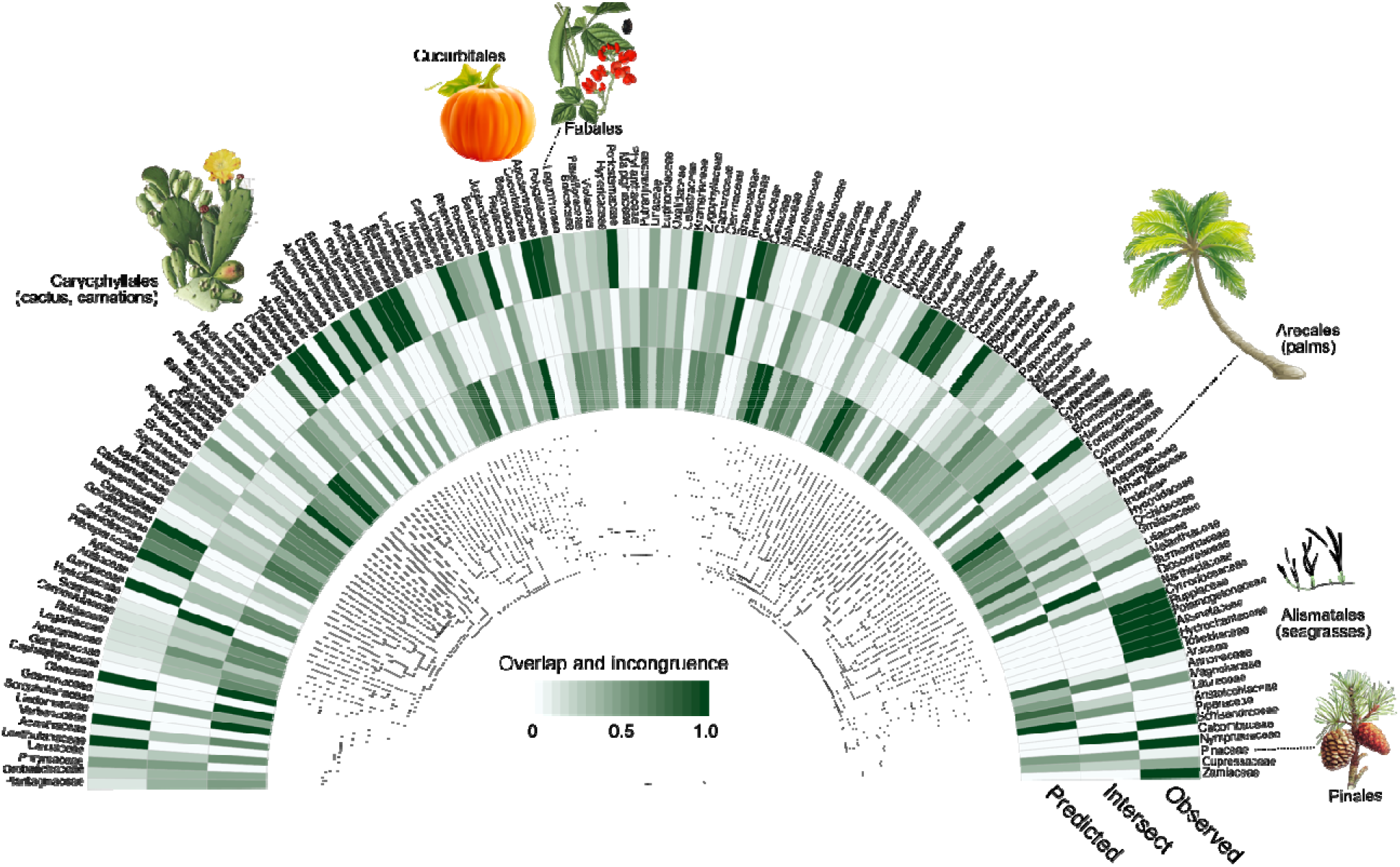
Phylogenetic representations of observed versus predicted geographic distributions of vascular plants in Texas USA. Iconic plant clades are indicated with images. The phylogenetic tree is summarized to family level to aid visualization.

### 2.2 Design and Development of a Mobile-based Application of GreenMaps

When incorporated in a mobile app with the locations setting set to TRUE, GreenMaps can provide users with three possible outcomes for each query: “present” (when the predicted species actually occur in an area), “absent” (when no matches to the predicted species are found in the area), and “rare/introduced” when the species observed in an area are different to the predicted ones. Volunteers can also edit the input occurrence point maps by removing outliers or erroneous records such as points corresponding to big cities, botanical gardens or herbarium buildings, or by manually entering geographic coordinates of occurrence points (as a CSV file), if a species is validated to occur in its predicted range. User input will be checked and when expert-validated will be used to generate new maps that incorporate these changes. Because the goal of GreenMaps is to generate high quality range maps that can facilitate dynamic analyses in macroecology and complement efforts of major existing data hubs, information collected through this tool will be saved in Darwin Core format and continuously synchronized with existing repositories such as the GBIF and iNaturalist.

## 3.0 Assessment of Plant Distributions in Space and Plant Diversification on Mountains

The products from GreenMaps will not only address the Wallacean shortfall for plants, but offer exciting opportunities for biodiversity research, ecosystem monitoring and conservation prioritization. For instance, to understand how plant diversity is distributed in space, I used the preliminary result to map hotspots of species weighted endemism (Crisp et al. 2001; Laffan and Crisp 2003)—species richness inversely weighted by species ranges—by varying the analysis along global, regional and local scales at country level (**Fig. 5**). In a global setting, the key plant endemism hotspots are chiefly located in the tropics and include Mesoamerica, Andes, Amazonia, southern Africa, Madagascar, Himalaya-Hengduan mountains, and southeast Asia (**Fig. 5a**). At the continental scale, hotspots of species endemism were less spatially clumped and more dispersed into new locations including Western Europe, Southeast Asia, and Southwestern Australia (**Fig. 5b**). In parallel, some regions which emerged hotspots at the global scale including the Atlantic forest of Brazil and Madagascar disappeared in the continental scale analysis (**Fig. 5b**). At the national scale, spatial patterns of species endemism became more widespread across countries, clustering more at the political borders of countries (**Fig. 5c**). Thus, these data can provide relevant natural history information at the evaluated scale (global, continental, country or biome), highlighting regions where conservation measures will have the most impact. As a next step, the detailed range maps can be paired with a supertree for all the vascular species derived using the function *phylobuilder* from our new R package bioregion (Daru and Schliep 2019), to explore phylogenetic endemism (PE). PE is the phylogenetic equivalent of species endemism, and measures the degree to which a substantial proportion of phylogenetic diversity is restricted to an area (Rosauer et al. 2009). Although both WE and PE capture different facets of endemism and are increasingly considered crucial for conservation prioritization (Daru et al. 2019), PE is less sensitive to changes in taxonomic knowledge and captures additional information that can facilitate more informed conservation decisions.

**Fig. 5.**
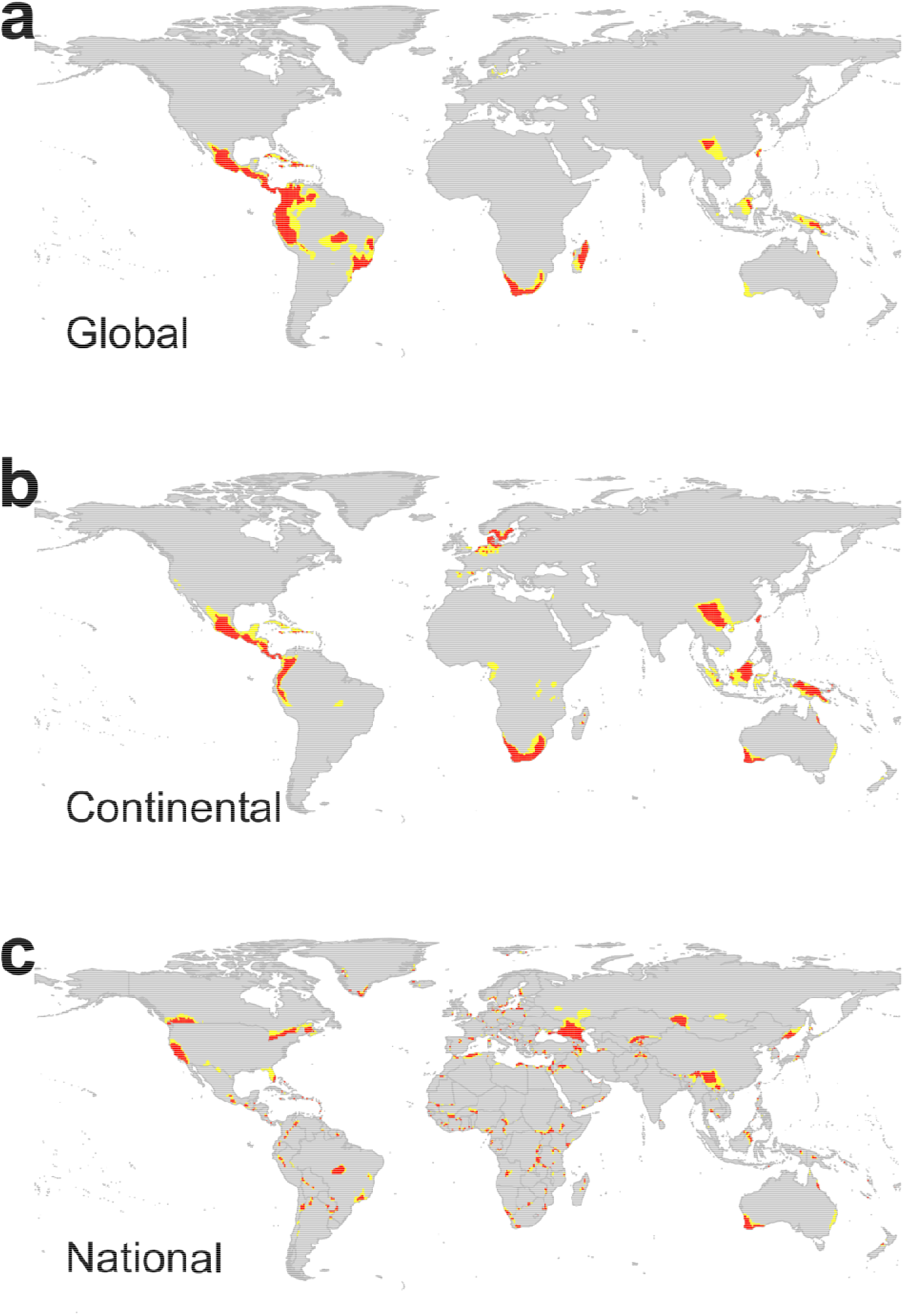
Variation in spatial extent of hotspots of species weighted endemism for vascular plants of the world across successive spatial extents (national, continental, and global). Hotspots are defined as the grid cells with the highest 2.5% of species richness (indicated in red), and 5% hotspots in yellow.

A global distribution of plants can provide insights into the tempo and mode of plant diversification in high elevations. Mountain ranges harbour disproportionately rich and unique biodiversity (Hoorn et al. 2013; Quintero and Jetz 2018), but face larger threats, particularly because the serve as buffers for species against global climate change (Sandel et al. 2011; González-Varo et al. 2017). Several explanations for the rich biodiversity found in mountains propose that orogeny creates conditions favouring rapid *in situ* speciation of resident lineages (Hoorn et al. 2013; Wen et al. 2014; Favre et al. 2015; Schwery et al. 2015; Hughes 2016; Lagomarsino et al. 2016), however, global assessment of mountain plant diversity is lacking. A comprehensive global geographic map of plants can provide insights into the processes underlying the assembly and maintenance of mountain biodiversity and their evolutionary origins. For example, we can use detailed distribution maps and phylogenetic relationships to test the relationship between elevation, species richness and speciation rate for plants across the major mountain systems of the world that represent theatres of spectacular evolutionary events such as the Drakensberg in southern Africa, Eastern Arc, Hengduan, Himalayas, and Andes. If species richness gradients are controlled by differences in speciation, extinction and migration – perhaps because species at higher elevations tend to have lower speciation rates or higher extinction rates (Kozak and Wiens 2010) – then we should observe a decrease in diversification rates and lower species richness with elevation. I also hypothesize that mountain age (time since uplift), seasonality, primary productivity and climate change will explain substantial variation in the species richness and diversification rates of mountain species. The modeled range maps can be combined with detailed phylogenetic relationships (e.g. Zanne et al. 2014; Smith and Brown 2018) to estimate species richness along elevation for plants at a global scale. Speciation rates can be estimated using BAMM (Rabosky et al. 2017), a Bayesian framework for reconstructing complex evolutionary dynamics from phylogenetic trees. To the extent that speciation rates exhibit a signal, the effect of other variables including mountain age and climate can be tested.

Other outstanding questions remain: 1) What are the effects of reduced area and increased isolation of habitats?, 2) Where are the hotspots of plant diversity, endangerment and endemism?, 3) Where will alien exotic plants spread to in future?, 4) Where should we locate protected area networks and conservation corridors to safeguard plant diversity? 5) Where will native plant species disperse to under alternative scenarios of climate change? 6) How have anthropogenic activities e.g. habitat conversion and urbanization change the geography of nature?, and 7) Which geographic locations to target for undiscovered species?

## 4.0 Incentivizing citizen science contribution and outreach

Two common challenges of citizen science campaigns include attracting and retaining a wide range of volunteers from all over the world, and ensuring that good quality data is generated (Fritz et al. 2009). The GreenMaps campaign will use a combination of educational feedback (regarding natural history, rarity, endemism), and quizzes, to incentivize volunteer participation.

Educational feedback constitutes a key element in quality control (Fritz et al. 2017). GreenMaps archive data is designed with an in-built system that allows participants to learn about centers of high conservation potential near their locality in real-time e.g. centers of endemism (**Fig. 5**) or identify regions of sampling bias in botanical collecting (see **Fig. 1**). Areas of endemism, where species with exceptionally small geographic ranges concentrate, are targets for conservation because they capture facets of biodiversity not represented elsewhere (Myers et al. 2000; Ceballos and Ehrlich 2006; Rosauer et al. 2009). From any 0.5 degree grid cell on the planet, volunteers can get instant feedback on the number of plant species predicted to occur versus actual observations around them (e.g. see **Fig. 3**) relative to the national, continental or global flora. Similarly, users can get instant feedback on geographic biases in botanical sampling density. This approach is vital to ensure that volunteers do not waste time in areas that are oversampled, but make effective use of their efforts, and allows them to see where botanical sampling data alone disagree with the predicted range maps of plant species. In the short-term, these approaches can lead to stronger appreciation of data as scientific output.

Mobile learning is another way to incentivize student participation in the GreenMaps campaign as it has the potential to extend education beyond the traditional classroom (Hlodan 2010). In the short-term, biology quizzes and game elements such as those used for most computer games e.g. *Pokemon*, could be added to GreenMaps to make participation more engaging. The long-term goals will be that such approach can improve student appreciation of natural history and can potentially lead to more interest in considering a career in biodiversity science.

Ensuring good quality data is crucial. I propose to implement a feature in a similar way to *Amazon* star ratings that allows entries to be monitored by volunteers through votes and comments by anyone who disagrees with them or filter out records with low star ratings (see **Fig. 1**). The data generated will be publically available and serve as a useful resource for science (e.g. biogeography) and conservation managers.

## 5.0 Concluding remarks

By integrating expert maps, point occurrence records, species distribution models and citizen science validation, this framework will enable wider use of species geographic distribution information particularly for less charismatic groups, such as plants with very sparse and biased occurrence data, but which play key crucial roles for diversity at other trophic levels as well as human health and well-being. Modelling allows predictions of species geographic distributions based on environmental conditions and expert-delineated range maps. Cloud-scale computing can significantly make the predictions faster particularly when the number of species is large, spanning more than 194,000 species, and the number of point occurrences per species are too few for estimating robust species distribution models. In turn, citizen science input is recorded along with ancillary data and geolocations for future use to generate new and improve global maps of plant diversity. Thus, GreenMaps combines strengths from heterogeneous data sources to improve predictions of plant species geographic distributions in a statistical way, a significant step toward addressing the Wallacean shortfall and can facilitate dynamic biodiversity analyses.

## Acknowledgements

I specially thank Klaus Schliep, Peter C. le Roux, Carolyn Weaver, Israel Loera, Tammy L. Elliott, Emily Meineke for valuable comments and discussion during the early formation of this paper. My appreciation goes to Texas A&M University-Corpus Christi for logistic support.

